# Effectiveness of the protected areas on the Mornington Peninsula for the common resident shorebird species using citizen science data

**DOI:** 10.1101/2021.09.28.462220

**Authors:** Udani A. Wijewardhana, Pragalathan Apputhurai, Madawa Jayawardana, Denny Meyer

## Abstract

In the absence of comprehensive survey data this study used citizen science bird counts, extracted from the Atlas of Living Australia, to assess which species benefit most from protected areas. This was done by fitting temporal models using the Integrated Laplace Approximation (INLA) method.

The trends for five resident shorebird species were compared to the Australian Pied Oystercatcher, with significantly steeper upward trends identified for the Black-fronted Dotterel, Red-capped Dotterel and Red-kneed Dotterel. Steeper upward trends were observed in protected than unprotected areas for the Black-fronted Dotterel, Masked Lapwing and Red-kneed Dotterel.

This work suggests that, with some limitations, statistical models can be used with citizen science data for monitoring the persistence of resident shorebirds and for investigating factors that are impacting these data. The results for the Dotterel species in protected areas are particularly encouraging.

## Introduction

In this paper we consider changes in the numbers of resident shorebirds on the Mornington Peninsula, situated close to Melbourne, Australia. This peninsula runs from the mainland to the Bass Strait with Western Port Bay on its eastern side and Port Phillip Bay on its western side. Volunteers of Parks Victoria and BirdLife Australia are involved in the monitoring and protection of nesting shorebirds on the Mornington Peninsula (Schmidt, 2019). Their primary focus is in providing protection for Hooded Plovers (Cuttriss et al., 2015; Dowling and Weston, 1999), an endangered local shorebird found in this area. Their other activities include the management of other threatened species, habitat protection and enhancement, and revegetation. Reserves are also managed to minimize the impact of bushfire.

We used citizen science data to track changes in the reported number of sightings for six resident shorebird species between 2010 and 2019 in response to the creation of protected areas. As explained below these species vary in terms of their vulnerabilities, with the creation of protected areas proving to be a successful method for addressing some of these vulnerabilities. In addition, we discuss the reasons for the use of citizen science data in this study and the use of advanced statistical modelling procedures for analysing these data.

### Species Vulnerabilities

Shorebirds make up about 10% of Australia’s bird species (Department of Agriculture, Water and the Environment, 2020). Most Australian shorebirds are in decline due to risk factors such as loss of habitat and human activities. Australian shorebirds occupy and breed in a wide variety of habitats and wetland types. Some species such as the Sooty Oystercatcher inhabits both sandy and rocky ocean beaches and their associated mudflats. Australian Pied Oystercatchers favour tidal mudflats, sandbanks and sandy ocean beaches. Red-capped Dotterels inhabit broad sandy or muddy shores of tidal or inland saline wetlands. Masked Lapwings use a wide range of open habitats, including pasture, sports ovals and mown lawns as well as associated wetlands including freshwater wetlands and tidal mudflats, usually preferring a combination of grassland and wetland. The Black-fronted and Red-kneed Dotterels inhabit freshwater wetlands (Marchant & Higgins 1993; Higgins & Davies 1996).

Tidal flats in Australia have been damaged by or lost to reclamation, altered water regimes, pollution, sea-level rise and weed invasion (Oldland and BirdLife Australia, 2009; The Great Barrier Reef Marine Park Authority, 2012). Other factors affecting species vulnerability on the Mornington Peninsula are oil spills, dog walkers, introduced rodents, avian predators such as ravens and silver gulls, reptilian predators and weed such as marram grass, sea spurge and sea wheatgrass (Maguire et al., 2019). Damage of habitat reduces access to food as well as breeding success, with this effect felt more keenly by bird species that nest on the ground as most shorebirds do, with birds spending less time providing care for fledglings when stressed. For these reasons it is hypothesised that seven common resident Mornington Peninsula shorebird species which all nest on the ground, Red-capped Plover (*Charadrius ruficapillus*), Red-kneed Dotterel (*Erythrogonys cinctus*), Black-fronted Dotterel (*Elseyornis melanops*), Masked Lapwing (*Vanellus miles*), Australian Pied Oystercatcher (*Haematopus longirostris*) and Sooty Oystercatcher (*Haematopus fuliginosus*), will have shown declines in abundance in recent years.

Interestingly the Bird Observation and Conservation Australia (BOCA) (now BirdLife Australia) Westernport surveys which commenced in late 1973 and were conducted at least thrice-yearly since then, provide some support for this hypothesis. Their survey methodology (Loyn et al. 1994; 2001) focused on the extensive intertidal mudflats of Western Port, with results from these surveys published quite recently by Loyn et al. (2018). Of the 39 species analysed in the 43-year survey period (1974-2017), 22 species showed declines, including the locally breeding Masked Lapwing. However, it was noted that some species had increased, including the Australian Pied Oystercatcher (many of which breed on French Island, where foxes are absent).

### Protected Areas

Protected areas are a key strategy for preserving biodiversity, but few studies have evaluated the effectiveness of this strategy. However, Cazalis et al. (2020) have used citizen science data from the eBird platform (eBird.org, 2021) to show that creating protected areas is an effective strategy for retaining species of conservation concern. This conclusion was based on studies of eight tropical forest biodiversity hotspots in Asia, Africa and the Americas, where biodiversity was particularly threatened, and data was particularly scarce. Our study of protected areas is a much less ambitious project, considering only one section of an important Australian biodiversity hotspot, the Western Port Biosphere Reserve.

Figure 1 shows the protected areas of the Mornington Peninsula that contribute most to biodiversity conservation based on an analysis of remnant native vegetation cover and quality, landscape context and threatened species habitat obtained from Parks Victoria. These areas will benefit the most from actions that are targeted to enhance biodiversity (Schmidt, 2019). As proposed by Diamond (1975), birds are better protected if reserves are larger, round or square rather than long and thin, with multiple reserves linked by steppingstones or corridors, and containing at least one large reserve to ensure that local extinctions in small reserves can be replenished. The network of reserves shown in Figure 1 exhibits some of these characteristics in some areas. Based on the above it is therefore hypothesised that reductions in the number of shorebirds counts will be greater outside than inside these protected areas.

**Figure 1.**
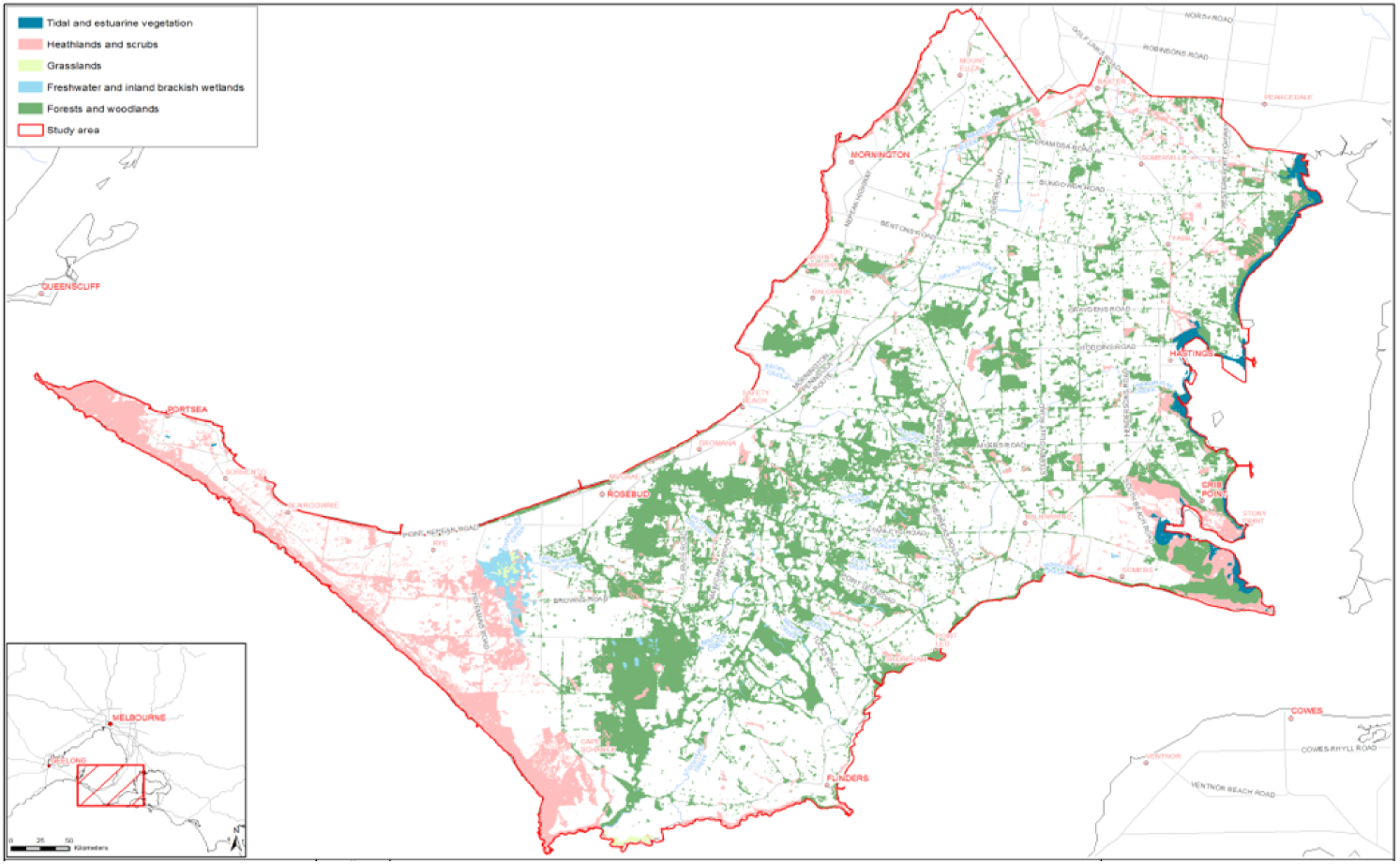
The protected areas that contribute most to biodiversity conservation on Mornington Peninsula Shire

**Figure 2.**
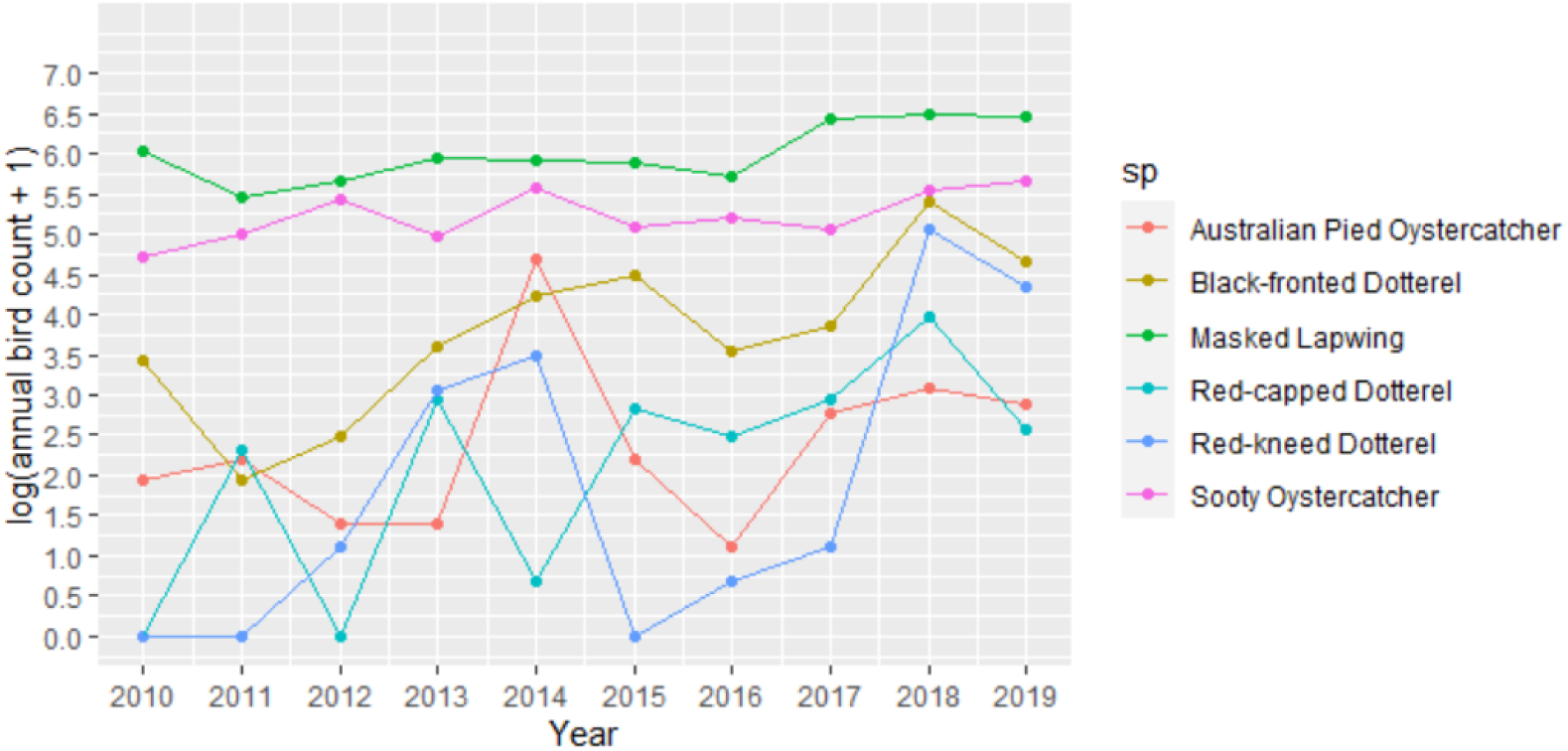
Annual Species Counts for the Mornington Peninsula from 2010-2019

### Citizen Science Data

Citizen science data plays an increasingly important role in the conservation domain. Continually updated, reliable and comparable biodiversity data is necessary to implement international conservation policy (Schmeller et al., 2015). Citizen science involving non-professionals and professionals as contributors can provide an intensive source of species observation data. In traditional citizen science studies, citizens are trained to collect samples using a standardised method to ensure high data quality. For instance, as a consequence of the continuous, long running Western Port survey, the significance of Western Port for birds has become widely recognised. As a result, it has become a key monitoring site for Palaearctic and Australasian shorebirds (Dennett and Loyn, 2009). Conservation actions using citizen science data range from research and monitoring to conservation planning, including tangible conservation actions such as site and habitat management, species management and habitat protection informing law and policy. The people involved benefit from hands-on learning experiences (Bonney and Dhondt, 1997) and improved environmental awareness (Gustafsson et al., 2015). The preparation and participation of professional volunteers helps raise community awareness of environmental issues while also contributing to the collection of data that would otherwise be too expensive to acquire (Hansen et. al., 2015).

The data presented in this study were obtained from a validated citizen science database (Nicholls, 2011), namely the Atlas of Living Australia (ALA). The ALA receives support from the Australian Government through the National Collaborative Research Infrastructure Strategy (NCRIS) and is hosted by the Commonwealth Scientific and Industrial Research Organisation (CSIRO). ALA data management supports the collection and sharing of data with documented quality parameters, with systems developed to implement the process of data collection, digitisation, validation, cleaning and access. The ALA creates filters for their data allowing the removal of duplicate entries, spatially suspect records and records for which scientific naming of species is not clear (Nicholls, 2011). In this study we have only used the validated ALA data and have ignored data classed as sensitive.

However, there are valid concerns about the detection bias existing in citizen science data, as explained by Iknayanm et al. (2013), and this is a limitation of studies using citizen science data such as this one. In addition, there has been increasing citizen science effort in recent years which means that trends in absolute numbers of sightings must be treated with caution. The Annual Report 2018-19 for the Port Phillip and Western Port Catchment Management Authority provides an example of the power of citizen science for monitoring the persistence of birdlife in the vicinity of Melbourne, Australia. Between the 2005-06 and 2016 analyses the number of wildlife sightings increased from 437,845 to over 3 million dues to a proliferation of citizen science survey programs. However, the large amount of data provided by citizen scientists means that more advanced statistical modelling techniques can be applied, as described below.

## Methodology

### Statistical Models for Bird Abundance

The development of species distribution models has benefited biodiversity conservation through their linkage of science to policy and decision processes. These models have evolved to provide scenarios for future landscapes based on known and projected environmental parameters. However, spatial or temporal scales can confound inference about changes in species observation data when used to draw conclusions about potential impacts on a different scale. Our data is confined to a relatively small spatial area over a short period of time (2010-2019), and for this reason we only consider temporal models. The Poisson distribution (Cameron and Trivedi, 1998) is assumed for the annual bird counts and significance is detected when credibility intervals do not contain zero.

The Integrated Laplace Approximation (INLA) method is an approximation tool for fitting Bayesian models for species abundance. INLA is an alternative robust method for the traditional Markov Chain Monte Carlo (MCMC) based Bayesian analyses (Paul et al., 2010). The key advantages of INLA are the ease with which complex models can be created and modified, without the need to write complex code, and the speed at which inference can be done, even for temporal problems with hundreds of thousands of observations (Sarul, 2015).

In this study we have used ALA citizen science data and INLA modelling in order to investigate the health of resident shorebird species on the Mornington Peninsula. In this study, we investigate the effects of climate and conservation reserves on shorebird abundance for seven shorebird species. It is likely that the effects of these variables will be different for each shorebird species, dependent on their specific vulnerabilities. The effects of climate are of particular importance because of the projected warming expected in the future due to climate change, while the creation of new conservation reserves are part of Australia’s national response for conserving our biodiversity and protecting our ecosystems (Department of Agriculture, Water and the Environment, 2020).

We have divided our analysis into two main sections. First, we tested whether there are significant differences in the trends for different species, to determine which species are more/less vulnerable. This is done using the entire data set using the species variable with seven categories. Then, we tested the effects of protected areas for each of the species. This is done for each species using a binary variable to indicate the level of protection for the location of each recorded sighting.

### Data Sources and study area

Records of annual data for resident shorebirds for the Mornington Peninsula from 2010 to 2019 were obtained from the Atlas of Living Australia database. The resident shorebird species included in this study are: Black-fronted Dotterel, Masked Lapwing, Red-capped Dotterel, Red-kneed Dotterel, Australian Pied Oystercatcher and Sooty Oystercatcher. The Atlas of Living Australia database gathers basic data on bird abundance and distribution at a variety of temporal scales. Much of this geocoded observational data were gathered during systematic surveys made by trained volunteers or qualified biologists (“Atlas of Living Australia”, 2021; Nicholls, 2010).

The Mornington Peninsula is a unique place when it comes to biodiversity. It is home to a wide range of plants and animals, including species of regional, state, national and international significance. The Mornington Peninsula is built of complex geological formations, resulting in diverse landforms and habitat types. The major habitat types on the Peninsula include central hills, waterways, wetlands, north central plains, sandy beaches and dunes, cliffs and headlands, rocky shores, mudflats, saltmarsh, mangrove swamps and estuaries. Approximately 10% of the Peninsula’s land is protected within parks or reserves, including a national park, a state park, state conservation reserves, local bushland reserves and foreshore coastal reserves. These remnant areas of bushland provide an important refuge for the peninsula’s diverse range of plants and animals.

The Mornington Peninsula forms part of the Western Port Biosphere Reserve (WPBR), which covers five Local Government Areas around Western Port Bay and carries out projects to test ways of balancing conservation with development. The Western Port wetlands are included in the Ramsar List of Wetlands of International Importance and are the primary reason why the United Nations Education, Scientific and Cultural Organisation (UNESCO) declared the Western Port catchment as one of only four active biosphere reserves in Australia, and one of only 701 in the world (Schmidt, 2019). As explained by Lopoukhine et al. (2012), the Western Port Ramsar Site was designated as a Wetland of International Importance under the Convention on Wetlands of International Importance (Ramsar Convention) in 1982 (Kellog et al., 2010). It regularly supports a high diversity and large numbers of waterbirds (Kellog Brown & Root 2010).

The Mornington Peninsula National Park is the largest reserve on the peninsula with inland and coastal components. Mornington Peninsula National Park is the most visited National Park in Victoria, with intensively used recreation nodes at Portsea, Sorrento and Cape Schanck (Schmidt, 2019). Because it is located close to residential areas and is such a popular holiday destination, the Mornington Peninsula is subject to a wide range of threats and pressures for shorebird species. The shorebirds in this area are especially threatened by human disturbance, recreational activities and predation. However, the Mornington Peninsula Shire undertakes a range of conservation programs to protect and enhance biodiversity and reduce the impact of threats posed by environmental weeds, pest animals and habitat loss.

### Statistical Analysis

We fitted temporal models for the number of annual sightings recorded for the above resident shorebirds on the Mornington Peninsula between 2010-2019. A Bayesian hierarchical modelling approach was used to conveniently account for parameter uncertainty and potential temporal dependence.

We have considered temporal models with Poisson and Negative Binomial distributions (Cameron et al., 1998). In most cases better fit, measured using the lowest Deviance Information Criterion (DIC) and Watanabe-Akaike information criterion (WAIC) was obtained for the Poisson distribution as defined below, with *λ* > 0 defined as the mean of this distribution and *k* the number of sightings in a single year.

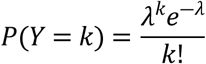

As described below we define a general hierarchical model for annual total shorebird counts (*Y*) in terms of categorical predictor variables with the annual trend.

For a categorical variable, such as species or level of protection, the general hierarchical model can be described as follows, where *Y*_*tj*_ is the annual shorebird counts for a specific category (*j*) in a particular year *(t*) and *f*(*y*_*tj*_|*λ* _*tj*_) is the Poisson distribution defined above with relevant mean parameters (*λ*_*tj*_).

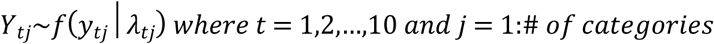

We include the categorical variables in the model by way of dummy variables (υ), with one of the categories chosen as the reference category. The interaction between year and these dummy variables tells us about the difference in slope between these categories and the reference category and allows a test of significant.

For species *j* the general link function for the expected annual total shorebird counts is given below with *δ* ’s included as species parameters. The Australian Pied Oystercatcher was chosen as the reference category in this analysis because this species showed the least change in the number of annual sightings over time (See Figure 3).

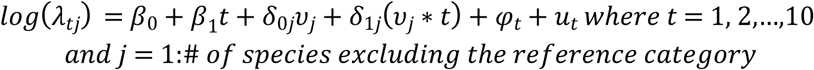

**Figure 3.**
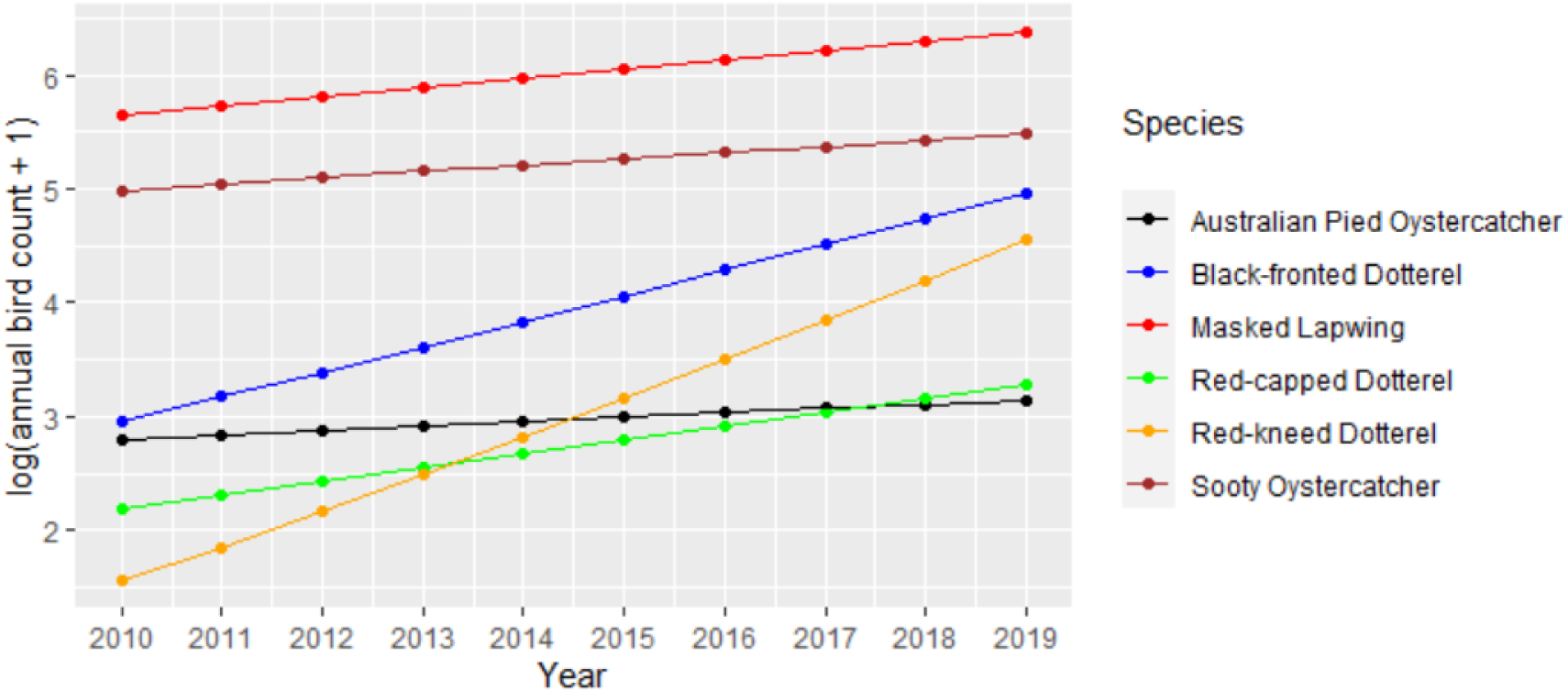
Model predictions of log transformed trend models for annual species sightings for the Mornington Peninsula

Depending on whether or not a sighting occurred in a protected area the general link function for the expected annual total shorebird counts is given below with *δ* ‘s included as protection parameters (See Figure 5).

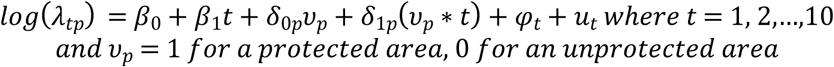

**Figure 5.**
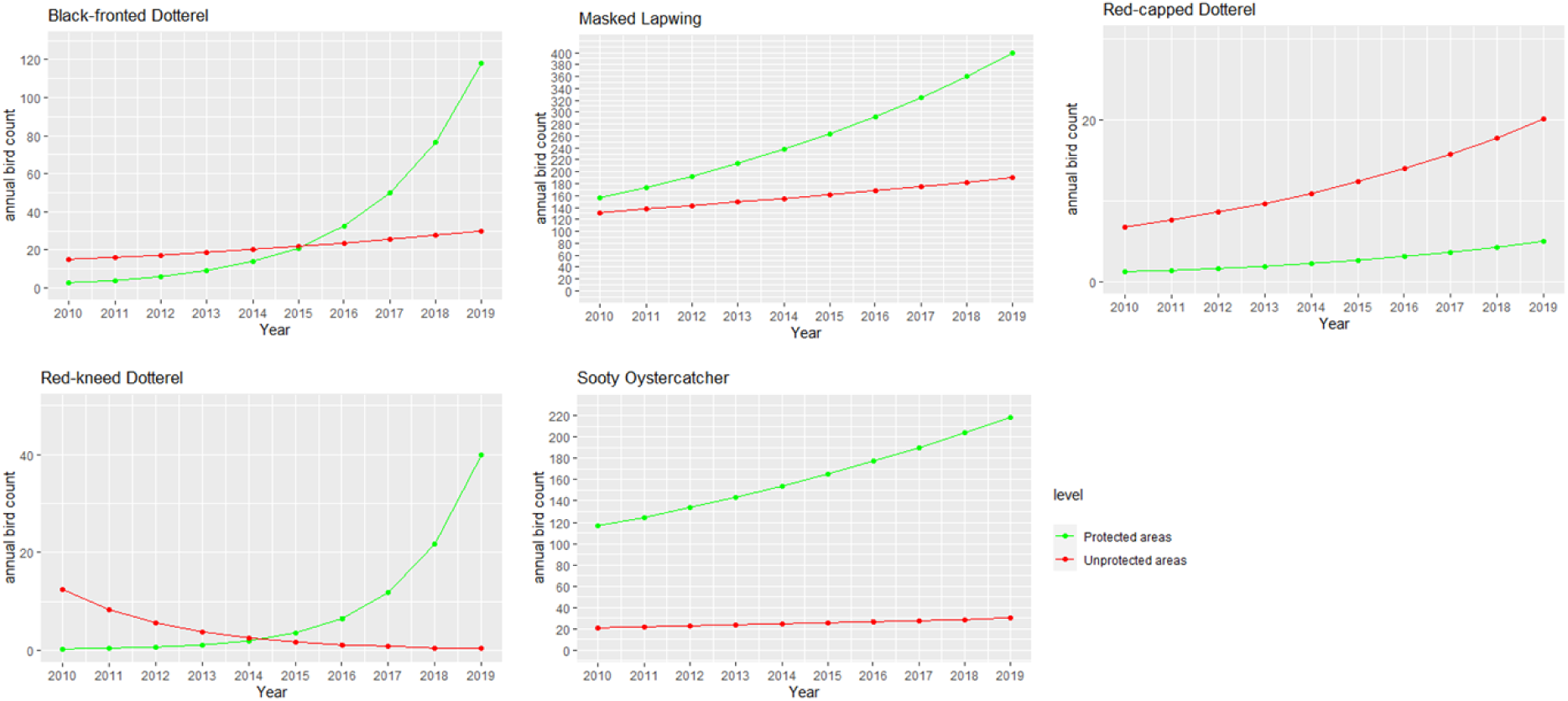
Model predictions of trends for the local shorebird species significant for protected and unprotected areas

As before *φ*_*t*_ allows for a temporal random effect with first-order autoregressive dependence and the random error term is represented by *u*_*t*_. All these models were implemented using the R-INLA (Rue et al., 2018) approach in R 3.6.1 (R core team, 2021). We have implemented a shiny app for annual temporal models using R-INLA which can be accessed from https://github.com/uwijewardhana/UDMTA.

## Results

### Descriptive Statistics

Table 1 shows that the Atlas of Living Australia data was mainly sourced from eBird together with the estimated annual growth rates in the number of sightings and the percentage of sightings recorded in protected areas for each species. Figure 2 illustrates the changes in annual sightings recorded for the seven resident shorebirds over the period 2010-2019 on a logarithmic scale.

**Table 1.** Bird Counts extracted from Atlas of Living Australian by Data Source (2010-2019)

| Species | BirdLife | eBird | Victorian Biodiversity Atlas | Total Counts | Estimated Growth per annum (%) 2010-2019 | % sightings in protected areas |
| --- | --- | --- | --- | --- | --- | --- |
| Australian Pied Oystercatcher | 3 | 84 | 103 | 190 | 4.7 | 10% |
| Black-fronted Dotterel | 110 | 519 | 15 | 644 | 26.6 | 54% |
| Masked Lapwing | 1715 | 2328 | 213 | 4256 | 9.2 | 58% |
| Red-capped Dotterel | 8 | 129 |  | 137 | 14.6 | 18% |
| Red-kneed Dotterel | 8 | 279 |  | 287 | 44.5 | 88% |
| Sooty Oystercatcher | 293 | 1632 | 1 | 1926 | 6.4 | 87% |

### Species Effects

Table 2 provides the results for a model testing whether there was significant difference in the trends for the various species. In this model the Australian Pied Oystercatcher has been chosen as the reference level because it showed the least change in the number of annual sightings and showed the weakest trend over time. When compared to the Australian Pied Oystercatcher there is a significantly stronger trend for the Black-fronted Dotterel, Red-capped Dotterel and Red-kneed Dotterel with estimated growth rates of 19%, 9% and 32% per annum. This significance is indicated by the credibility intervals for their *δ*_1*j*_ estimates which do not contain zero. Figure 3 illustrates the above trends on a logarithmic scale.

**Table 2.** Fixed effects for species trends using the Australian Pied Oystercatcher as the reference species (significant effects bolded)

| Coefficients | Mean | SD | 95% Credibility Interval |  |
| --- | --- | --- | --- | --- |
|  |  |  | 0.025quant | 0.975quant |
| Intercept |  |  |  |  |
| Australian Pied Oystercatcher - reference ($\beta_0$) | 2.701 | 0.293 | 2.147 | 3.32 |
| Black-fronted Dotterel ($\delta_{01}$) | -0.03 | 0.203 | -0.424 | 0.371 |
| Masked Lapwing ($\delta_{02}$) | 2.854 | 0.167 | 2.535 | 3.19 |
| Red-capped Dotterel ($\delta_{03}$) | -0.759 | 0.328 | -1.415 | -0.127 |
| Red-kneed Dotterel ($\delta_{04}$) | -1.734 | 0.323 | -2.38 | -1.111 |
| Sooty Oystercatcher ($\delta_{05}$) | 2.223 | 0.171 | 1.896 | 2.566 |
| Trend by Year |  |  |  |  |
| Australian Pied Oystercatcher - reference ($\beta_1$) | 0.04 | 0.044 | -0.053 | 0.124 |
| Black-fronted Dotterel ($\delta_{11}$) | 0.188 | 0.029 | 0.131 | 0.245 |
| Masked Lapwing ($\delta_{12}$) | 0.042 | 0.025 | -0.008 | 0.091 |
| Red-capped Dotterel ($\delta_{13}$) | 0.091 | 0.045 | 0.004 | 0.179 |
| Red-kneed Dotterel ($\delta_{14}$) | 0.317 | 0.041 | 0.238 | 0.398 |
| Sooty Oystercatcher ($\delta_{15}$) | 0.015 | 0.026 | -0.036 | 0.066 |

### Effects of Protected Areas

There are many ongoing conservation programs on the Mornington Peninsula, especially for the Hooded Plover. Our results in Table 3 show that the protected areas are particularly important for the resident shorebirds including locally threatened Sooty Oystercatcher. However, although the bird counts for the Black-fronted, Red-capped and Red-kneed Dotterels were initially low, the results show that the latest trend is starting to reverse this effect with significantly greater growth seen in protected areas. Figure 4 illustrates these upward trends while also emphasising the obvious growth in Masked Lapwing numbers.

**Table 3.** Fixed effects of protected level models using the unprotected areas as the reference level with significant effects bolded.

| Species | Coefficients | Mean | SD | 0.025quant | 0.975quant |
| --- | --- | --- | --- | --- | --- |
| Australian Pied Oystercatcher | Intercept | 1.589 | 1.034 | -0.529 | 3.676 |
|  | year | 0.107 | 0.16 | -0.22 | 0.433 |
|  | protected | -3.519 | 2.9 | -9.771 | 1.612 |
|  | year:protected | 0.192 | 0.325 | -0.392 | 0.885 |
| Black-fronted Dotterel | Intercept | 2.613 | 0.808 | 1.151 | 4.452 |
|  | year | 0.078 | 0.121 | -0.199 | 0.296 |
|  | **protected** | **-2.166** | **0.293** | **-2.753** | **-1.603** |
|  | **year:protected** | **0.354** | **0.038** | **0.28** | **0.431** |
| Masked Lapwing | Intercept | 4.841 | 0.286 | 4.35 | 5.517 |
|  | year | 0.04 | 0.042 | -0.056 | 0.117 |
|  | protected | 0.111 | 0.073 | -0.032 | 0.254 |
|  | **year:protected** | **0.063** | **0.011** | **0.042** | **0.084** |
| Red-capped Dotterel | Intercept | 1.786 | 0.677 | 0.397 | 3.102 |
|  | year | 0.121 | 0.092 | -0.068 | 0.301 |
|  | **protected** | **-1.759** | **0.919** | **-3.655** | **-0.047** |
|  | year:protected | 0.036 | 0.117 | -0.187 | 0.272 |
| Red-kneed Dotterel | Intercept | 2.918 | 3.517 | -4.796 | 9.547 |
|  | year | -0.405 | 0.484 | -1.375 | 0.602 |
|  | **protected** | **-5.325** | **1.192** | **-7.965** | **-3.294** |
|  | **year:protected** | **1.015** | **0.234** | **0.624** | **1.539** |
| Sooty Oystercatcher | Intercept | 2.991 | 0.25 | 2.452 | 3.448 |
|  | year | 0.041 | 0.039 | -0.031 | 0.125 |
|  | **protected** | **1.695** | **0.153** | **1.4** | **1.999** |
|  | year:protected | 0.03 | 0.023 | -0.017 | 0.076 |

## Discussion

In this study we have tried to determine the effects of species, climate and protected areas for the numbers of the common locally-breeding shorebird species found on the Mornington Peninsula. This has been done using Atlas of Living Australia data and Poisson temporal models, fitted using the INLA method. We wanted to know if the rate at which the reported numbers are changing over time differs according to species and protected areas. Our findings raise several important issues for conservation.

Australian Pied Oystercatcher has increased significantly in Western Port over the past 40 years (Hansen et. al., 2015), however our study suggests only a 4.7% annual increase on the adjacent Mornington Peninsula in the period 2010-2019. Instead, contrary to expectation our results suggest that there has been a significant upward trend for the Black-fronted Dotterel, Red-capped Dotterel and Red-kneed Dotterel when compared to Australian Pied Oystercatcher. The increase in observer numbers explains the increase in citizen science data effort over the years. Possible reasons for the increase in Dotterel counts could be the result of a growing interest in these birds. For example, “The friends of the Hooded Plover Mornington Peninsula” started dedicated data collection for the Red-capped Dotterel in 2010. This may have resulted in increased survey effort for these birds in recent years.

Declines in population were reported for both the Masked Lapwing and the Red-capped Dotterel up to 2009 (Hansen, Menkhorst, Loyn, 2011). However, the 9.2% annual growth rate for ALA sightings for the Masked Lapwing observed in this study and the 14.6% annual growth rate for the Red-capped Dotterel suggest that these trends have since reversed, or did not apply to the different habitats sampled in our study.

The most vulnerable habitats for resident shorebirds on the Peninsula are coastal, especially for shorebirds with restricted distributions such as the Sooty Oystercatcher. In our models for protected areas, we found that there was a significant beneficial effect for the locally threatened Sooty Oystercatcher. The higher counts for the Sooty Oystercatcher can perhaps be explained by their locally threatened status, making the focus of regular surveys and the location of protected areas.

In addition, this study has found that the percentage increase for the numbers of Black-fronted Dotterel, Red-kneed Dotterel and Red-capped Dotterel has been stronger in protected than unprotected areas on Mornington Peninsula since 2010, and the same has been true for the Masked Lapwing. Contributing factor are likely to include the control of introduced predators (Red Fox (*Vulpes vulpes)*, Cat (*Felis catus)*, Black Rat (*Rattus rattus*,) domestic Dog (C*anis familiaris*) in protected areas. The core habitat for Red-kneed and Black-fronted Dotterels is inland wetlands. The study period (2010 - 2019) coincided with severe inland drought, with the inland becoming increasingly dry through the study period. This may have been the cause of the increase in the Mornington Peninsula area as inland wetlands dried up. The increasing trends may also mean that the monitoring of the local Dotterels is more pervasive than for other birds in protected areas, producing the higher counts observed in these areas. However, for Pied Oystercatcher, there is no evidence to suggest that protected areas are successful for increasing the number of shorebird counts.

Australian Pied Oystercatcher, Masked Lapwing and Red-capped Plover make extensive use of tidal mudflats among other habitats. Only peripheral data on species that inhabit freshwater habitats (Black-fronted and Red-kneed Dotterels), rocky or sandy shores (Sooty Oystercatcher) existed previously. On Mornington Peninsula, most of the protected areas relevant to shorebirds are on scenic beaches, tidal mudflats, reserves which has freshwater habitats and sandy shores. This difference in habitat between the Mornington Peninsula and previously reported studies may be the main reason for some of the changes noted above. The ALA sighting data considered in this study do not confirm earlier positive trends reported by Hansen et al. (2015) for the Australian Pied Oystercatcher. Also, it was previously reported that Masked Lapwing numbers were declining, and this study suggests that this is no longer the case. However, the previous study focused on a different habitat (intertidal mudflats in the Western Port survey) as opposed to the broad range of habitats sound on the Mornington Peninsula. In addition, there have been many changes since 2009 including the creation of additional protected areas and fox eradication programs which may have impacted the trends observed in this study.

The use of unstructured citizen science data in this study is an obvious limitation. As discussed above, the nature of these data and their focus on threatened species makes some degree of bias inevitable. In particular, the ALA growth rates reported in this paper must be treated with caution because to some extent they reflect the increased effort made by citizen scientists. Another limitation of this study is that the analysis uses only annual data. We have used annual shorebird sightings in order to avoid seasonal bias in the data and to ensure adequate sample sizes for the INLA modelling, however, the shortness of the time series considered (only 10 years) makes this another limitation of our analysis.

## Conclusions

Temporal models fitted using INLA provide flexible and useful frameworks for modelling count data using statistical models. While citizen science data has its problems, especially for comparing the relative abundance of different species, there are clear advantages for using citizen science data for monitoring the abundance of resident shorebird species. For these species the quality of the citizen science data tends to be better, at least in terms of the survey data collected in protected data, and it would be difficult to obtain sufficient data from other sources. However, growing counts for other birds, such as Dotterels in protected areas, shows growing citizen science effort for less endangered species and this bodes well for future conservation efforts.

This study has suggested that the protected areas are beneficial for local shorebirds. This study has therefore suggested that statistical models may indeed have a role to play in the future, for addressing the challenges associated with monitoring the persistence of resident shorebirds and for investigating factors that are impacting these data.

## Data Availability Statement

The data for this study is provided as supplementary materials together with the R-Code used for the analysis.

## Acknowledgement

Port Phillip and Westernport Catchment Management Authority (PPWCMA) for their interest in this project. In addition, we would like to thank Parks Victoria to provide us the protected areas in Mornington Peninsula and the citizen scientists who collected the data and the Atlas of Living Australia for making the data available.

